# Protection of human ACE2 transgenic Syrian hamsters from SARS CoV-2 variants by human polyclonal IgG from hyper-immunized transchromosomic bovines

**DOI:** 10.1101/2021.07.26.453840

**Authors:** Theron Gilliland, Yanan Liu, Rong Li, Matthew Dunn, Emily Cottle, Yutaka Terada, Zachary Ryckman, Maria Alcorn, Shauna Vasilatos, Jeneveve Lundy, Deanna Larson, Hua Wu, Thomas Luke, Christoph Bausch, Kristi Egland, Eddie Sullivan, Zhongde Wang, William B. Klimstra

**Affiliations:** Center for Vaccine Research and Department of Immunology, University of Pittsburgh, Pittsburgh, PA 15261; Department of Animal Dairy, and Veterinary Sciences, Utah State University, Logan UT 84341, United States; SAb Biotherapeutics, Inc. Sioux Falls, SD 57104

**Author notes:** These authors contributed equally (Theron Gilliland, Yanan Liu, Rong Li, Matthew Dunn).

## Abstract

Pandemic SARS CoV-2 has been undergoing rapid evolution during spread throughout the world resulting in the emergence of many Spike protein variants, some of which appear to either evade antibody neutralization, transmit more efficiently, or potentially exhibit increased virulence. This raises significant concerns regarding the long-term efficacy of protection elicited after primary infection and/or from vaccines derived from single virus Spike (S) genotypes, as well as the efficacy of anti-S monoclonal antibody based therapeutics. Here, we used fully human polyclonal human IgG (SAB-185), derived from hyperimmunization of transchromosomic bovines with DNA plasmids encoding the SARS-CoV-2 Wa-1 strain S protein or purified ectodomain of S protein, to examine the neutralizing capacity of SAB-185 *in vitro* and the protective efficacy of passive SAB-185 antibody (Ab) transfer *in vivo*. The Ab preparation was tested for neutralization against five variant SARS-CoV-2 strains: Munich (Spike D614G), UK (B.1.1.7), Brazil (P.1) and SA (B.1.3.5) variants, and a variant isolated from a chronically infected immunocompromised patient (Spike Δ144-146). For the *in vivo* studies, we used a new human ACE2 (hACE2) transgenic Syrian hamster model that exhibits lethality after SARS-Cov-2 challenge and the Munich, UK, SA and Δ144-146 variants. SAB-185 neutralized each of the SARS-CoV-2 strains equivalently on Vero E6 cells, however, a control convalescent human serum sample was less effective at neutralizing the SA variant. In the hamster model, prophylactic SAB-185 treatment protected the hamsters from fatal disease and minimized clinical signs of infection. These results suggest that SAB-185 may be an effective treatment for patients infected with SARS CoV-2 variants.

## INTRODUCTION

SARS CoV-2 has spread worldwide during the previous year resulting in over 140 million cases and over 3 million deaths (WHO dashboard https://covid19.who.int)^1^. In late fall 2020, variant viruses were identified that exhibited altered infection, transmission and disease characteristics ^2^ reviewed in ^3-6^. These variants may reflect immune response escape mutants as well as mutants adapting to replication and transmission in normal or immunocompromised human populations ^7 8,9^, reviewed in ^10,11^. Of particular concern are variants with multiple changes in the Spike protein that is a primary target of acquired immune responses, since these viruses could exhibit increased resistance to Spike-targeted vaccines and immuno-therapeutics ^7,8,12,13^ reviewed in ^14-17^.

Genetically modified transchromosomic bovines (Tc-bovines) adaptively produce fully human polyclonal antibodies after exposure to environmental or vaccine antigens ^18-20^. After hyperimmunization, Tc-bovines produce high titer, fully human IgG (Tc-hIgG) that can be rapidly produced from their plasma ^21-23^. Tc-hIgGs have shown pre-clinical efficacy against Middle East Respiratory Syndrome coronavirus (MERS-CoV), Ebola and Venezuelan equine encephalitis viruses among others ^21-23^. Here, we tested the *in vitro* neutralizing capacity and *in vivo* protective efficacy of a Tc-Bovine derived immunoglobulin (SAB-185) derived human IgG preparation ^24^ against four SARS CoV-2 variants using a new human ACE2 receptor transgenic hamster model ^25^. Efficacy evaluation of variant strains is important as SAB-185 is currently being evaluated in an adaptive Phase 2/3 treatment efficacy trial in multiple countries as sponsored by the National Institute of Allergy and Infectious Diseases (https://clinicaltrials.gov/ct2/show/NCT04518410?term=sab-185&cond=covid&draw=2&rank=3).

## RESULTS

Production and purification of the SAB-185 human IgG preparation has been described^24^. Briefly, transchromosomic bovines were hyperimmunized two times with a DNA plasmid expressing the WA-1 S protein followed by three times with recombinant spike ectodomain produced and purified from insect cells. Plasma was collected on 8, 11 and 14 days post each booster from V3 to V5. Then the qualified V3, V4 and V5 plasma was pooled and subjected to cGMP purification for human IgG SAB-185 Lot 6 which was for use in neutralization and hamster protection studies. We initially tested the neutralization capacity of SAB-185 and a National Institute for Biological Standards and Control convalescent human serum (NIBSC 20/136) for each of the variant viruses by plaque assay on Vero E6 cells (Figure 1 A, B). SAB-185 neutralized each of the viruses similarly with PRNT_50_ values ranging from 1:5,223-1:27,785 (Figure 1B). The NIBSC effectively neutralized the Munich, UK, Δ144-146 and Brazil variants (PRNT_50_ range 1:1,079-1:6,219) but was significantly less effective at neutralizing the SA variant failing to provide 80% or 50% neutralization endpoints at a 1:320 dilution (Figure 1B).

**Figure 1.**
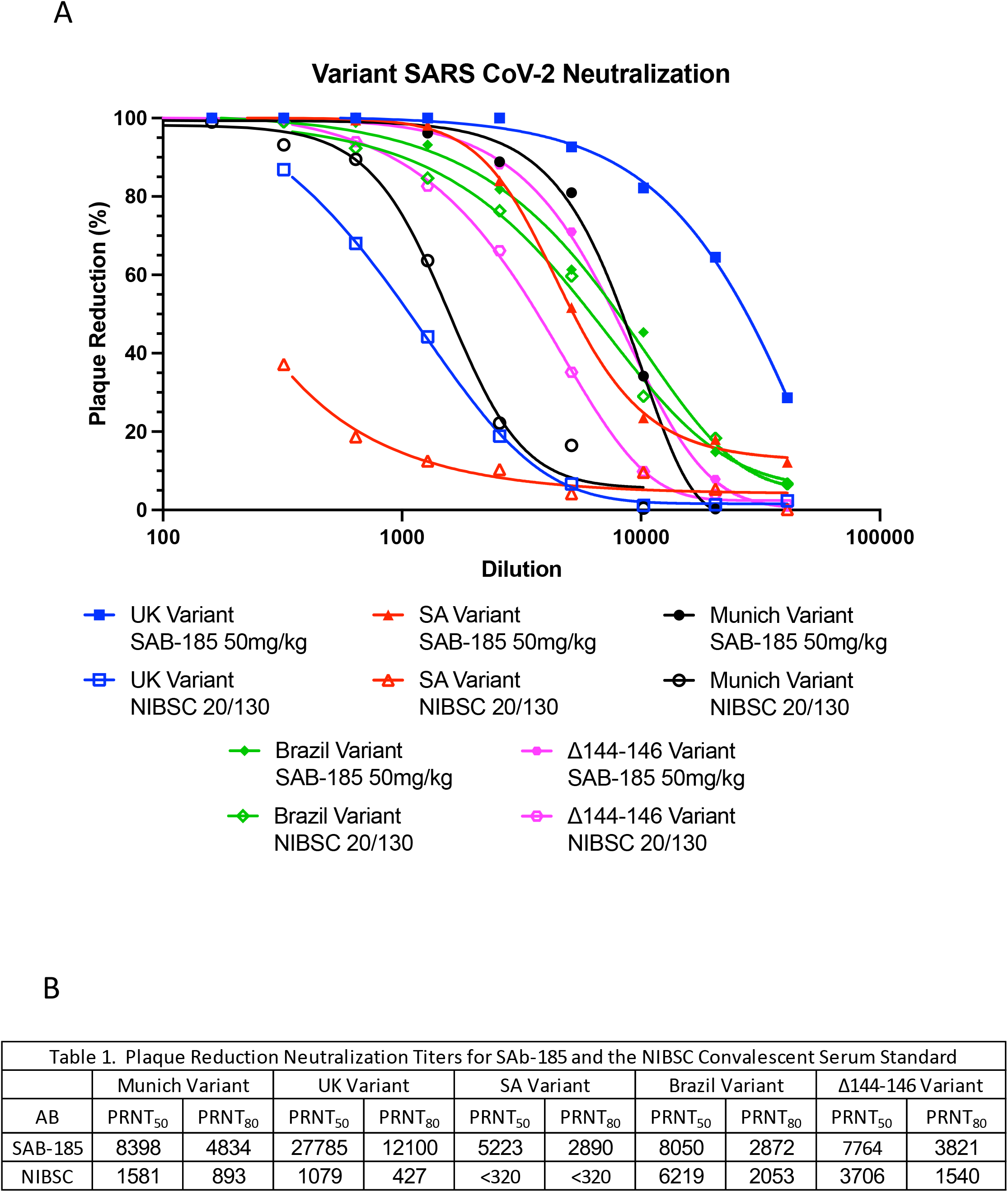
Neutralization of variant SARS CoV-2 isolates by SAB-185 and an NIBSC convalescent serum standard. A) SAB-185 was diluted to 1mg/ml in PBS and then diluted serially two-fold before reaction with viruses. The NIBSC convalescent serum standard was diluted 1:10 in PBS prior to serial two-fold dilution. Three independent dilutions of each sample were performed and these were replicated in two averaged wells per dilution. Data points represent averages of the three independent dilutions. B) PRNT_50_ and PRNT_80_ values were calculated as described in Materials and Methods.

The Syrian hamster is among the first rodents that have been successfully used as a model to study SARS CoV-2 infection, as hamsters are susceptible to wild type SARS CoV-2 without the needs of host adaptation and develop respiratory disease with some similarities to what are observed in COVID-19 patients ^24^. However, infected hamsters only develop limited clinical disease. While the hamster ACE2 protein has been shown to serve as a functional cell receptor for SARS CoV-2 infection, some amino acid residues critical for the recognition and binding by the SARS CoV-2 spike protein are not conserved between the hamster ACE2 and human ACE2 protein, which may diminish infectivity of SARS CoV-2 ^26,27^. To develop a highly susceptible hamster model mimicking severe infection in humans, we employed a piggyBac-mediated transgenic approach and generated multiple independent hACE2 transgenic hamster lines expressing the human ACE2 gene from the human cytokeratin 18 promoter ^28^. These hamster lines are highly susceptible to SARS CoV-2 infection *via* intranasal infection and develop respiratory disease and mortality, similar to that observed in severe COVID-19 patients ^25^. This model was used to test the protective efficacy of passive transfer of SAB-185.

Hamsters were injected intramuscularly in the gastrocnemius muscle with 50mg/kg SAB-185 followed 24 hours later by intratracheal challenge with 1000 Vero E6 plaque forming units of each variant virus. Controls were treated with PBS prior to challenge. In control groups, all of the female animals died between four and seven days (D) post-infection while male hamsters exhibited 50 to 75% mortality over a similar interval (Figure 2A). Times to death were not significantly different between the SARS CoV-2 variants (Figure 2A) (p>0.09 - p>1.0). SAB-185 treatment prior to infection completely protected hamsters from mortality regardless of hamster sex, which was significant for all viruses (Figure 2B-F).

**Figure 2.**
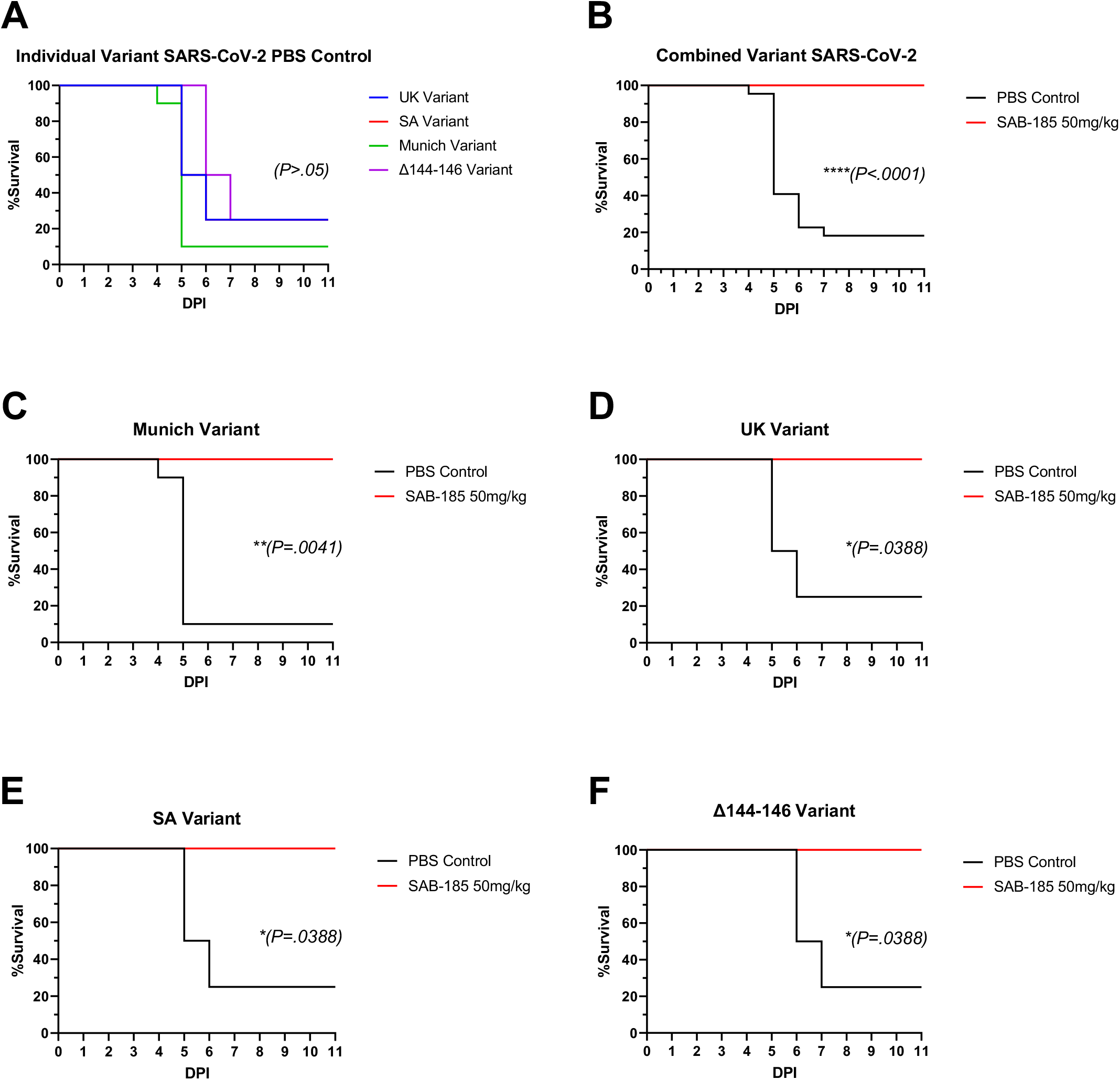
SAB-185 protection from mortality in hamsters challenged with four variant SARS CoV-2 isolates. Hamsters were administered SAB-185 or PBS intramuscularly and then challenged intratracheally 24 hours later with 1000 plaque forming units of each variant virus. Mortality for individual variant PBS controls (A) and for control and SAB-185 combined groups (B). Individual mortality data for Munich (C), UK (D), SA (E), and Δ144-146 (F) viruses. Mantel-Cox log-rank significance is indicated within each panel. * p<0.05, **p<0.01, ***p<0.005.

Weight loss was measured daily for all animals after challenge (Figure 3). Combined control group average weights began decreasing between D1 and D2 post challenge and were significantly different from SAB-185 treated groups animals on D4, 5 and 6 (Figure 3B). On D5, the last day on which all but one control animal were alive, Munich, and SA challenge groups were exhibited significantly lower weight loss than controls but the UK and Δ144-146 challenged animals did not (Figure 3E, G, I, K). All but one surviving mock-treated male animal lost weight over the course of the experiment suggesting that infection of that animal may have been very limited (Figure 3J). Only one SAB-185 treated animal (Δ144-146 group, Figure 3J) lost weight but this animal did not succumb to the infection. If mock survivors and the poorly infected SAB-185 animal (See virus genome titration section) were removed, D5 weight loss was significantly lower for Munich, UK and SA infected SAB-185 treated animals (Figure S1 D, E, F). The SAB-185 treated, Δ144-146 challenged animals also exhibited a non-significant trend towards less weight loss on D5 (Figure S1G)

**Figure 3.**
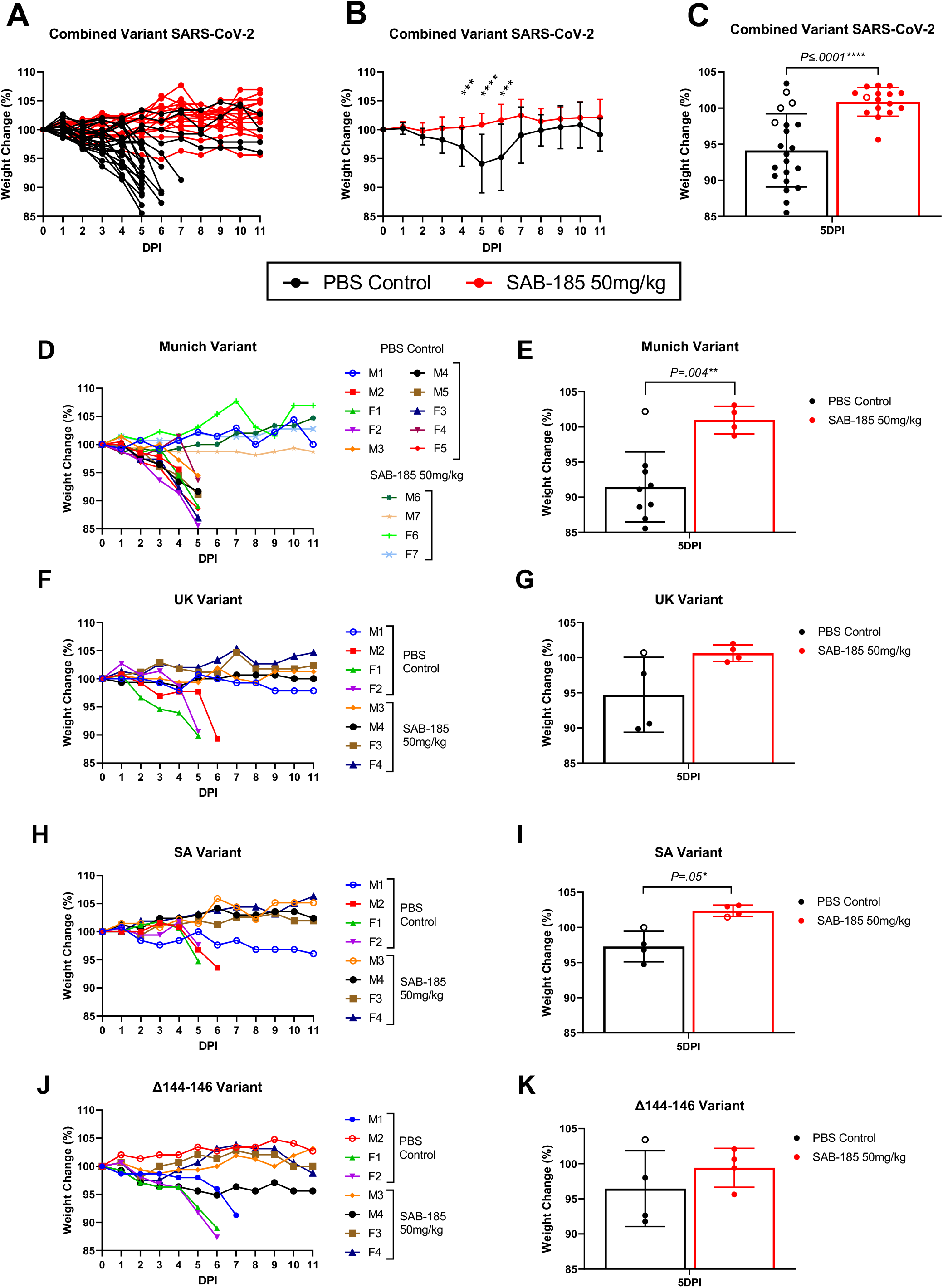
SAB-185 protection from weight loss in hamsters challenged with four variant SARS CoV-2 isolates. A) Weight loss for individual hamsters in all groups. B) Combined average weight loss data for SAB-185-treated and control hamsters. (C) Combined average weight loss data for SAB-185-treated and control hamsters on D5 (the last day most animals were alive) post challenge. (B). Individual weight loss data for Munich (D), UK (F), SA (H), and Δ144-146 (J) viruses. Combined average weight loss data for Munich (E), UK (G), SA (I) and Δ144-146 variants on D5 (the last day all animals were alive) post challenge. * p<0.05, **p<0.01, ***p<0.005. Open circles are surviving controls and the SAB-185 treated animal that exhibited delayed replication.

Clinical scoring data (hunching, ruffling, ataxia, seizures) for mock animals indicated general peaks of infection signs with all strains on D2 and D5 post challenge, suggesting similar disease profiles (Figure 4). Some SAB-185 treated animals also exhibited increased disease signs on these days but total scores were generally lower with the exception of the surviving male mock group animals. In combined data, clinical signs were significantly less *versus* controls in the SAB-185 group on D3-11 (Figure 4B) and highly significant on D5 and D6 (Figure 4B, C). Day 5 clinical scores were significantly different between the individual groups for the Munich and UK viruses but not SA or Δ144-146 (Figure 4E, G, I, K). However, if mock survivors and the poorly infected SAB-185 animal (See virus genome titration section) were removed, the SAB-15 treated Del 141-144 animals were also significantly different (Figure S2). Clinical signs were less severe in control animals challenged with the SA virus *versus* the other variants (Figure 4D, F, H, J) perhaps accounting for the smaller differences with this virus.

**Figure 4.**
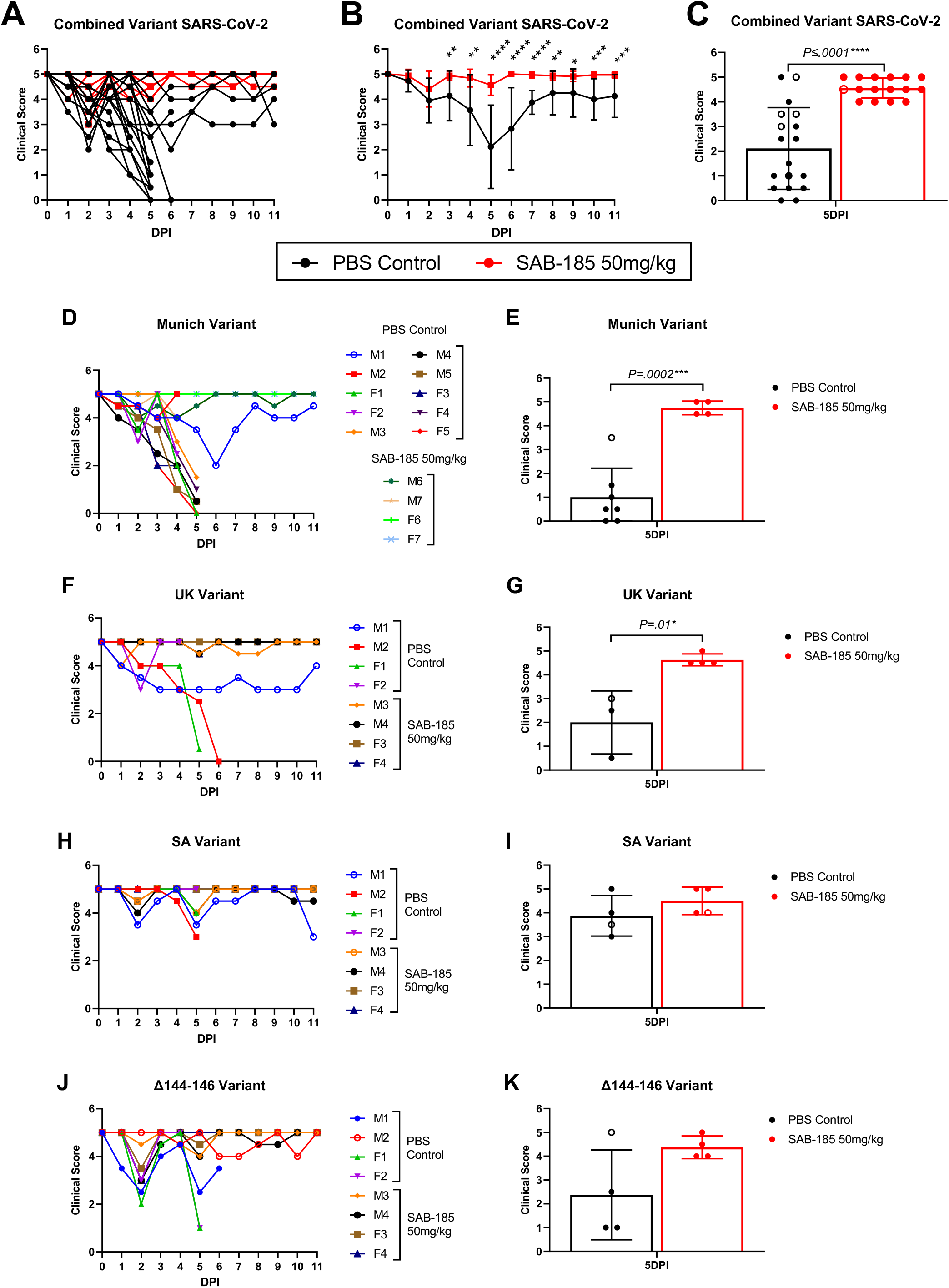
SAB-185 protection from clinical signs in hamsters challenged with four variant SARS CoV-2 isolates. Data is presented as the inverse of the clinical score sum values to be comparable to weight loss data. Each datum point represents an average of morning and afternoon observations. A) Clinical sign scoring for individual hamsters in all groups. Each datum point represents an average of morning and afternoon observations B) Combined clinical sign scoring data for SAB-185-treated and control hamsters. (C) Combined clinical sign scoring data for SAB-185-treated and control hamsters on D5 (the last day all animals were alive) post challenge. (B). Individual clinical sign scoring data for Munich (D), UK (F), SA (H), and Δ144-146 (J) viruses. Individual clinical sign scoring data for Munich (E), UK (G), SA (I) and Δ144-146 variants on D5 (the last day all animals were alive) post challenge. * p<0.05, **p<0.01, ***p<0.005. Open circles are surviving controls and the SAB-185 treated animal that exhibited delayed replication.

Oropharyngeal swabs were taken from animals and every other day through day 11 post challenge and assayed for virus genome equivalents (GE) by quantitative RT-PCR and plaque titration (Figure S3). GE titers in control animals were variable from undetectable (250 GE limit of detection) to >1×10^6^ GE/ml on day 1 post challenge suggesting similar initial replication among the variant strains (Figure S3A). Titers for most of the controls rose from D2 to D5 when most of them succumbed. Titers were highly variable between individual animals between combined control and SAB-185 treatment groups (Figure S3A) and none of the titers were significantly different (p>0.05) (Figure S3B, C). Notably, undetectable titers on D3-5 were associated with survival of the male animals in control groups. However, titers in all surviving animals had risen to detectable levels by D6 (Figure S3 A, D, F, H, J). Only one animal in the SAB-185 treated group exhibited this replication profile (SA variant challenge group Figure S3H), suggesting that only one of the SAB-185 animals might have survived without Ab treatment. As noted in weight loss and clinical sign sections above, if samples without titer on D3 to 5 were omitted from the analyses (including surviving controls and one SA challenged SAB-185 treated animal, Figure S4A-G), the combined data show significantly lower virus genome titers on D5 (Figure S4 C) with SAb-185 treatment. However, only the SA variant control group exhibited a significant difference with SAB-185 treatment. Oropharyngeal swab plaque titers for all hamsters were below the limit of detection of a plaque assay (25 PFU/ml) on D1, 3, 5, 7 and 11 post challenge suggesting that genomic RNA detected by the PCR assay was not associated with high levels of live virus.

## DISCUSSION

The current pandemic has yielded significant surprises for the scientific community especially concerning the rapid evolution of the SARS CoV-2 spike protein after infection of hundreds of millions of humans and the potential effect these adaptations could have on efficacy of vaccines and antibody based therapeutics. The current studies are the first to examine the efficacy of an antibody based therapeutic against multiple circulating SARS CoV-2 variants. Furthermore, we have demonstrated that a human IgG preparation derived from Tc-bovines hyperimmunized with antigens from a single SARS CoV-2 strain, can effectively neutralize and protect, with a single IM administration, against mortality and severe disease caused by multiple variant viruses. In contrast, a human convalescent serum standard was significantly less effective at neutralizing the SA virus strain *in vitro*.

The hACE2 transgenic hamsters used here also provide a new and relevant model of severe/fatal SARS CoV-2 disease that can be utilized to test vaccines and therapeutics^25^. Consistent with other studies of hamsters, other experimental animals and potentially humans ^29-32^, protection from severe disease was not closely associated with an effect on oropharyngeal swab titers; however, if control and SAb-185 treated animals that had delayed virus replication were removed, suppression of titers by SAb 185 was significant for combined challenge groups (p=0.02) on D5 post challenge. This contributes to the body of evidence that vaccination and/or Ab therapeutic treatment may only provide limited protection from virus infection and upper respiratory tract replication.

Reasons underlying the protective efficacy of SAB-185 *versus* multiple strains may include the hyper-immunization of TC-bovines, which may increase stimulation of polyclonal antibodies reactive with subdominant epitopes that are less likely to mutate during widespread human infection. Loss of reactivity of monoclonal antibodies that bind different epitopes in the S protein has been demonstrated clearly ^7,33,34^ and the broad reactivity provided by polyclonal Ab preparations may have an advantage in neutralizing and protecting against variants. In addition, since doses are scalable, more virus-reactive IgG can be administered with the Tc-bovine preparations than might be possible in passive transfer of human convalescent serum. Ultimately, the data in this report suggest that the human IgG preparations derived from hyperimmunized Tc-bovines may have efficacy in treating variant SARS CoV-2 infection of humans. Phase I clinical trials or SAB-185 have been recently completed and phase 2 clinical trials are ongoing.

---

**Figure S1.**
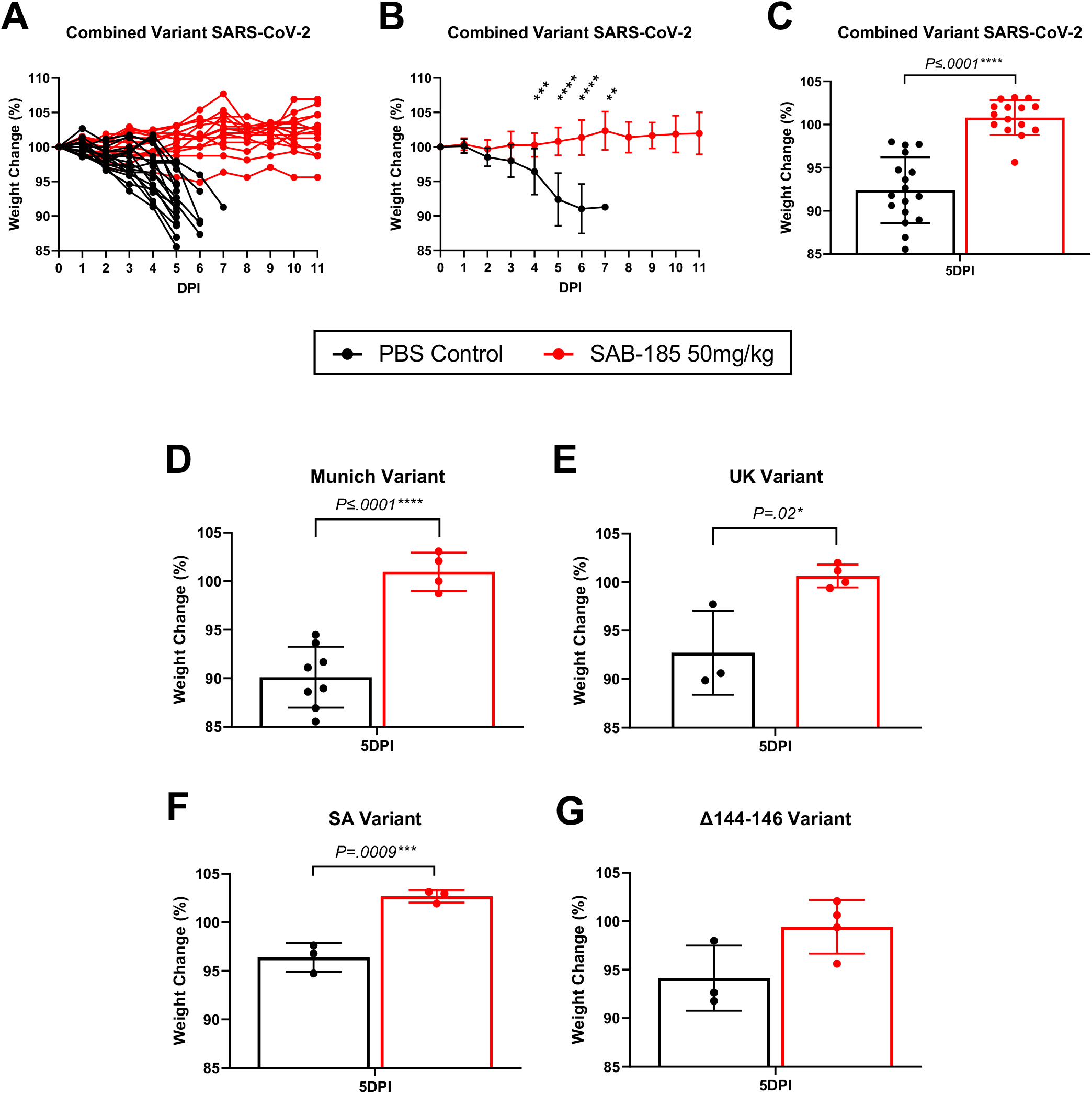
Weight loss data for control and SAB-185 treated hamsters with individual male animals that exhibited delayed replication removed. A) Weight loss data for individual hamsters in all groups. B) Combined weight loss data for SAB-185-treated and control hamsters. (C) Combined weight loss data for SAB-185-treated and control hamsters on D5 (the last day all animals were alive) post challenge. Individual averaged weight loss data on D5 for Munich (D), UK (E), SA (F), and Δ144-146 (G) viruses. * p<0.05, **p<0.01, ***p<0.005.

**Figure S2.**
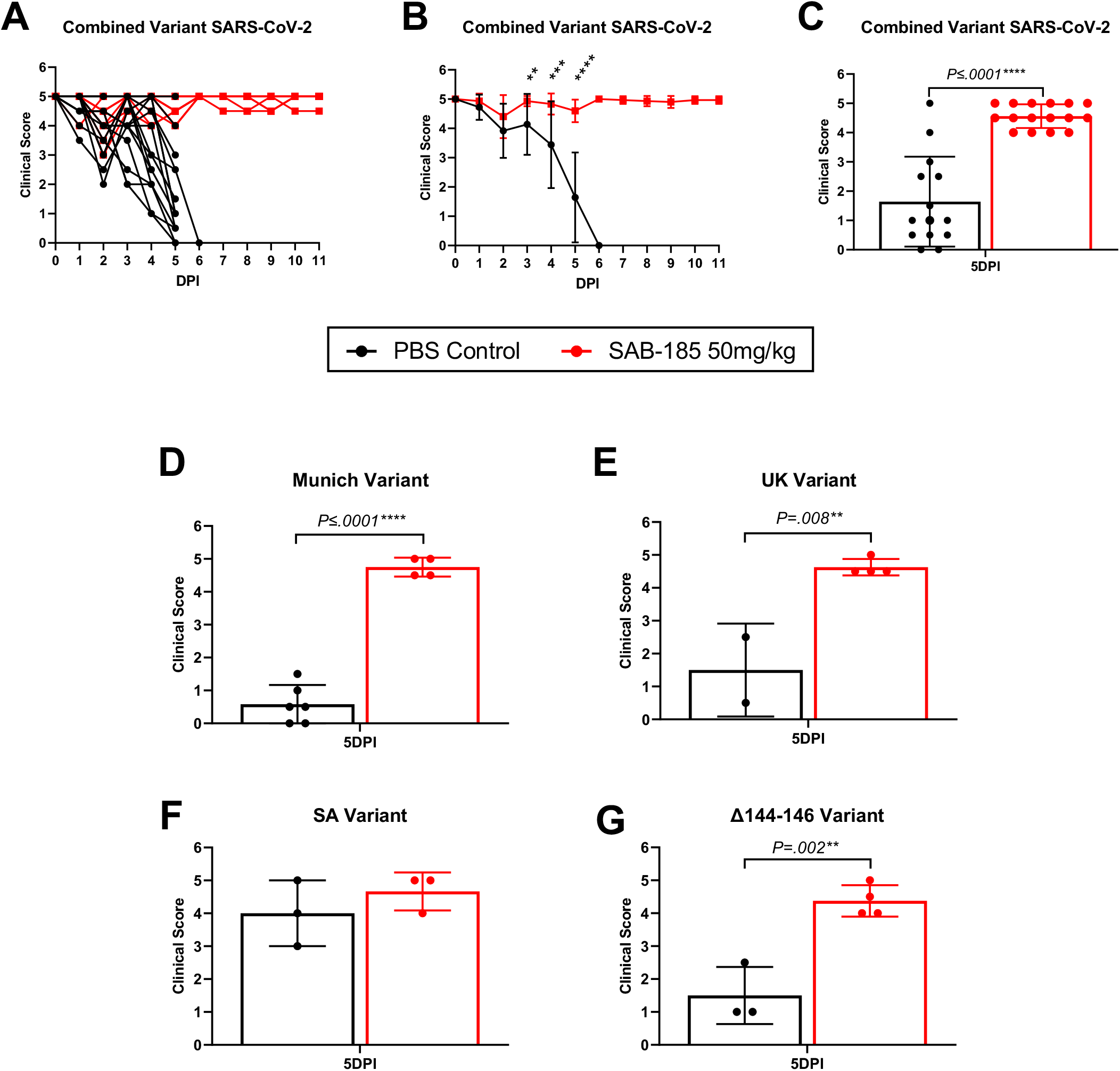
Clinical sign data for control and SAB-185 treated hamsters with individual male animals that exhibited delayed replication removed. Data is presented as the inverse of the clinical score sum values to be comparable to weight loss data. Each datum point represents an average of morning and afternoon observations. A) Clinical sign scoring for individual hamsters in all groups. B) Combined clinical sign scoring data for SAB-185-treated and control hamsters. (C) Combined clinical sign scoring data for SAB-185-treated and control hamsters on D5 (the last day all animals were alive) post challenge. Individual averaged clinical sign scoring data on D5 for Munich (D), UK (E), SA (F), and Δ144-146 (G) viruses. * p<0.05, **p<0.01, ***p<0.005.

**Figure S3.**
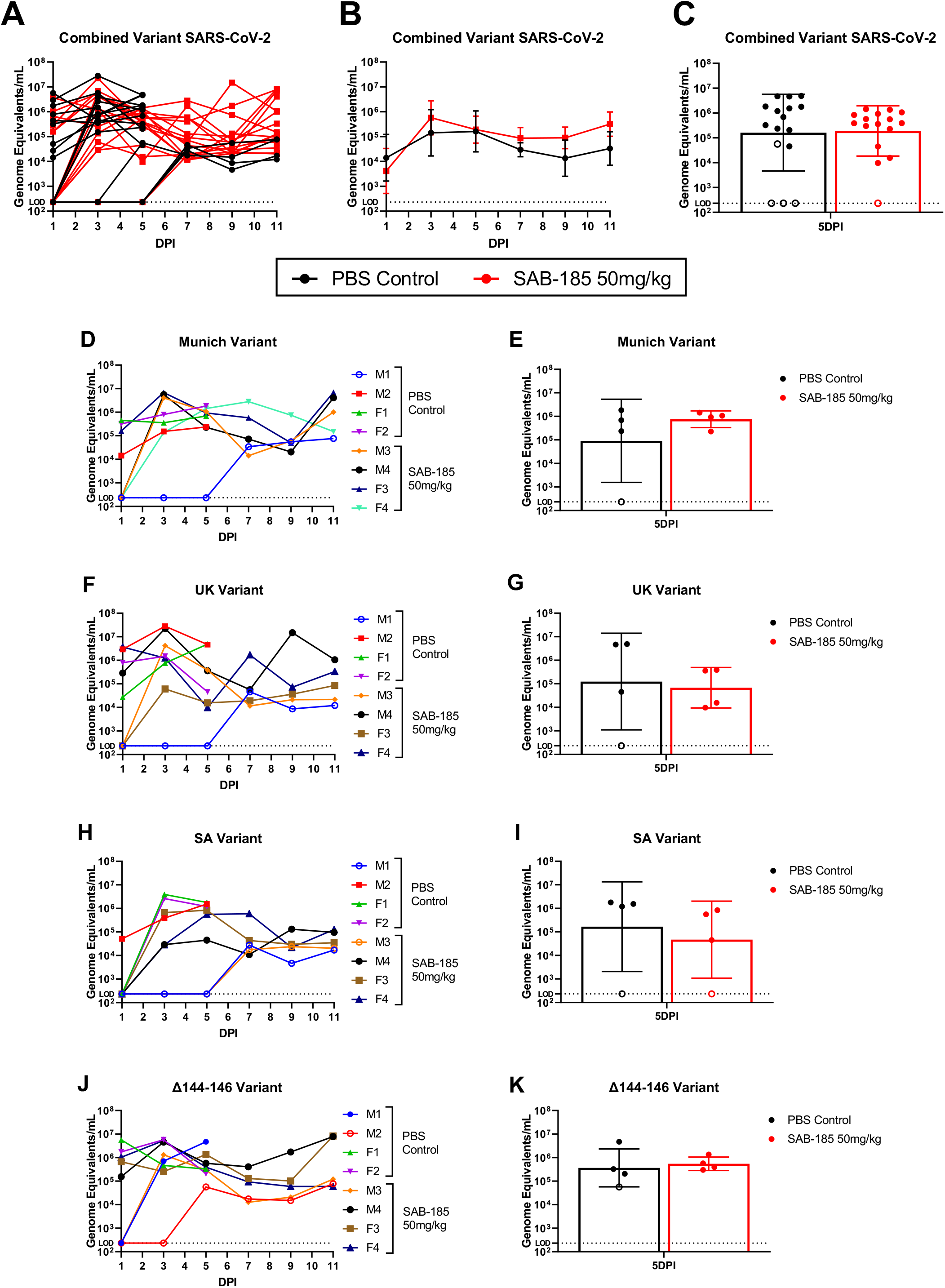
Quantitative RT-PCR for genome equivalents (GE) of SARS CoV-2 in oropharyngeal swabs for variant isolate-infected hamsters. Oropharyngeal swabs were collected days 1, 3, 5, 7, 9, and 11 post infection. Animals were sedated with isoflurane (3-5%) and a sterile swab was inserted into the oral cavity and upper trachea and then placed into virus diluent. Quantitative RT-PCR and plaque assays were performed on RNA purified from plasma and swab supernatants or directly with plaque assay as described in Materials and Methods. A) Quantitative RT-PCR GE data for individual hamsters in all groups. B) Combined quantitative RT-PCR GE data for SAB-185-treated and control hamsters. (C) Combined quantitative RT-PCR GE data for SAB-185-treated and control hamsters on day 5 (the last day all animals were alive) post challenge. (B). Individual RT-PCR GE data for Munich (D), UK (F), SA (H), and Δ144-146 (J) viruses. Combined RT-PCR GE data for Munich (E), UK (G), SA (I) and Δ144-146 variants on D5 (the last day all animals were alive) post challenge. *p<0.05, **p<0.01, ***p<0.005. The limit of detection was 250 GE. Open circles are surviving controls and the SAB-185 treated animal that exhibited delayed replication.

**Figure S4.**
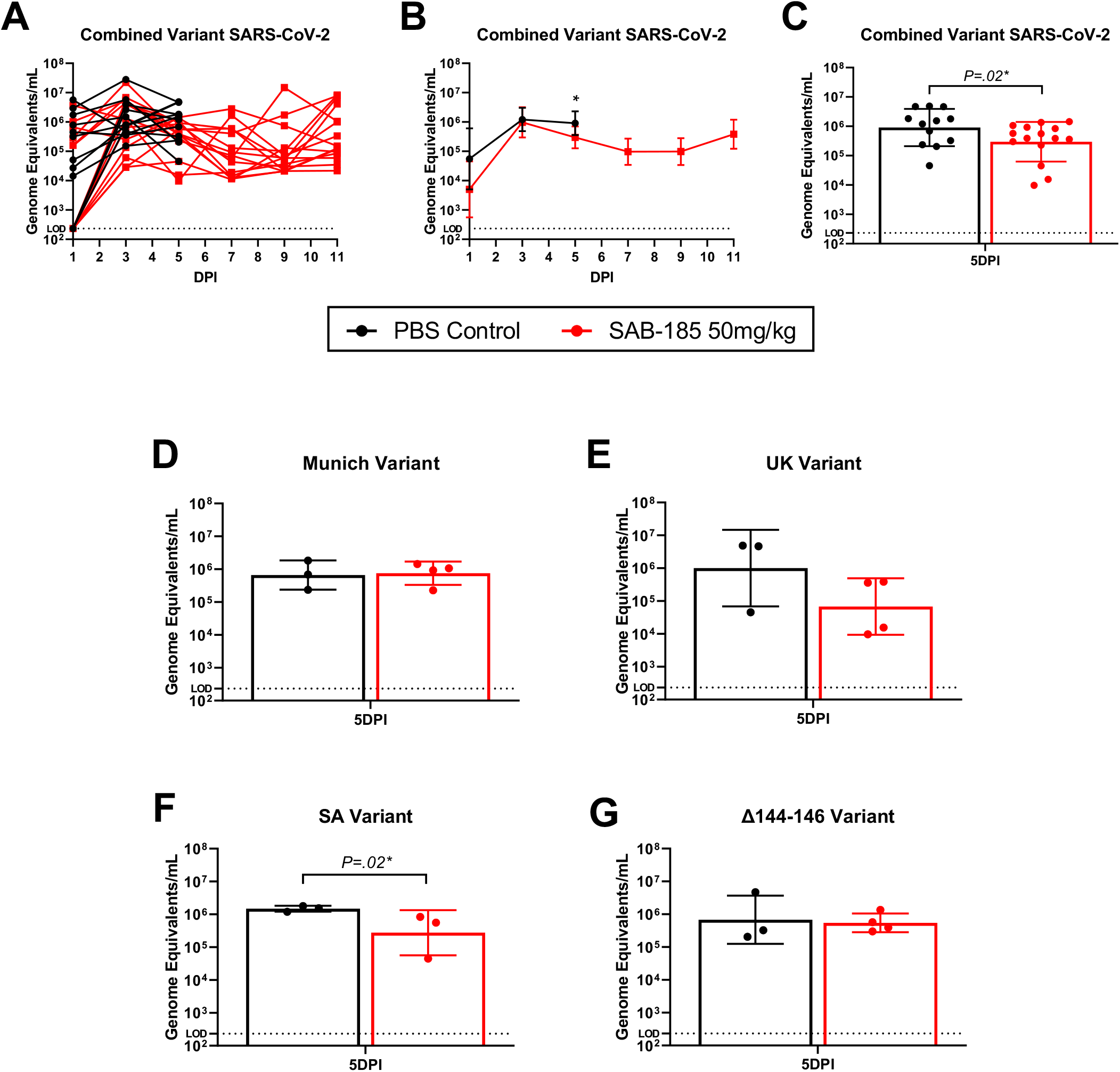
Quantitative RT-PCR for genome equivalents (GE) of SARS CoV-2 in oropharyngeal swabs for control and SAB-185 treated hamsters with individual male animals that exhibited delayed replication removed. A) GE data for individual hamsters in all groups. B) Combined GE data for SAB-185-treated and control hamsters. (C) Combined GE data for SAB-185-treated and control hamsters on D5 (the last day all animals were alive) post challenge. Individual averaged GE data on D5 for Munich (D), UK (E), SA (F), and Δ144-146 (G) viruses. *p<0.05, **p<0.01, ***p<0.005. The limit of detection was 250 GE.

## ACKNOWLEDGMENTS

SAB Biotherapeutics, Inc., is receiving support from the Department of Defense (DoD) Joint Program Executive Office for Chemical, Biological, Radiological, and Nuclear Defense (JPEO - CBRND) Joint Project Lead for Enabling Biotechnologies (JPL-EB), and from the Biomedical Advanced Research Development Authority (BARDA), part of the Assistant Secretary for Preparedness and Response (ASPR) at the U.S. Department of Health and Human Services, to develop SAB-185, a countermeasure to SARS-CoV-2 (Effort sponsored by the U.S. Government under Other Transaction number W15QKN-16-9-1002 between the Medical CBRN Defense Consortium (MCDC), and the Government). The US Government is authorized to reproduce and distribute reprints for Governmental purposes notwithstanding any copyright notation thereon. The views and conclusions contained herein are those of the authors and should not be interpreted as necessarily representing the official policies or endorsements, either expressed or implied, of the U.S. Government. HW, TL, CB, KE, and ES are employees of SAB Biotherapeutics and have financial interests. This work was supported by a contract from SAb Biotherapeutics, Inc., to the University of Pittsburgh (WK). MDHA was supported by an NIH/NIAID T32 grant (T32 AI049820).

## Author Contributions

TL, TG, HW, ZW, DR and WK designed the experiments. ZW, RL, DL and DR produced the hACE2 hamsters. TG, MD, EC, YT, ZR, MA, SV, JL, DR, and WK performed the experiments. TG, MD, TL, HW, ZW, CB, KE, ES and WK analyzed the data. WK and TG wrote the manuscript and TL, HW, ZW, DR and WK edited the manuscript.

## SUPPLEMENTARY MATERIALS AND METHODS

The Munich strain (containing D614G) was obtained and amplified in Vero E6 cells (ATCC CRL-1586) as described ^35^, while the UK and SA viruses were obtained from BEI Resources (BEI Resources, NIAID, NIH: SARS-Related Coronavirus 2, Isolate hCoV-19/England/204820464/2020, NR-54000 and BEI Resources, NIAID, NIH: SARS-Related Coronavirus 2, Isolate hCoV-19/South Africa/KRISP-K005325/2020, NR-54009, contributed by Alex Sigal and Tulio de Oliveira, respectively). The Δ144-146 virus (4 aa deletion in NTR RDR 2, a generous gift from Dr. Paul Duprex, University of Pittsburgh) was obtained from an immunocompromised patient as described and removes an epitope recognized by mAb 4A8 ^7,36^. The Munich (D614G; 4.0. x 10^6^ Vero E6 PFU/ml) and del 141-144 (5.0 x 10^5^ Vero E6 PFU/ml) viruses were isolates passaged three times in Vero E6 cells and shown to possess an intact S protein furin protease cleavage signal ^7,35^. The UK (4.0 x 10^6^ Vero E6 PFU/ml), SA (9.5 x 10^6^ Vero E6 PFU/ml) and Brazil (6.8 x 10^6^ Vero E6 PFU/ml) viruses were unamplified stocks from BEI Resources, diluted and used directly. Full genotypes of these stocks are available at https://www.beiresources.org.

Neutralization capacity of SAB-185 and the NIBSC convalescent serum control (NIBSC 20/136) was assayed by Vero E6 cell (grown in DMEM [Corning], 10% FBS [Atlanta Biologicals], 1 mM L Glutamine [Corning], 1x penicillin/streptomycin [Corning]) plaque assay. All Ab samples were heat inactivated by incubation at 56°C for 30 minutes. Viruses were diluted in OPTI-MEM (Gibco) with 2% FBS to approximately 200 PFU in 250 μl and reacted with an equal volume of serial two-fold dilutions of each antibody (in PBS) for 1 hour at 37°C followed by infection of Vero E6 monolayers for 1 hour at 37°C. A solution of 0.1% immunodiffusion agarose (MP Bio) in 2X Vero E6 growth medium was then added and plaques were developed at 37°C for 72 hours followed by removal of agarose, staining of cells with crystal violet (Fisher Scientific) and counting of plaques.

Equal numbers of male and female hamsters of 6-8 weeks of age were randomized by animal support staff and housed singly after receipt at the University of Pittsburgh. All hamster procedures were in accordance with AAALAC procedures and approved by the University pf Pittsburgh IACUC committee (protocol #IS00017405). Hamsters were challenged intratracheally with 50 μl of virus diluent (OPTIMEM medium; Gibco) containing 1000 Vero E6 cell plaque forming units of each virus. We observed a sex-based difference in mortality in this model with mock-treated females uniformly succumbing to infection with each tested virus but males exhibited approximately 50 to 100% mortality depending upon the experiment.

Hamsters were observed (∼30 seconds-1 minute) twice daily post challenge through day 12 (acute viral disease period). Weights were recorded once daily. After challenge with SARS CoV-2 hamsters exhibited ruffling of fur, hunching, ataxia, anorexia and weight loss. Hamsters losing >20% of starting body weight or exhibiting prolonged hunching/ataxia (>3 days) indicative of severe disease were euthanized. Animals were scored (0 for no, 1 for yes and a cumulative score totaled) for appearance of ruffled fur, hunching, increased respiratory rate, anorexia or lethargy.

One step quantitative RT-PCR was performed as previously described ^35^ for SARS CoV-2 RNA on blood taken day 3 post challenge and from oropharyngeal swabs taken at multiple times post challenge. Quantitative RT-PCR primer/probe sets (forward 5’ TTA CAA ACA TTG GCC GCA AA 3’, reverse 5’ GCG CGA CAT TCC GAA GAA 3’ probe: 5’ 6-FAM/ ACA ATT TGC CCC CAG CGC TTC AG /BHQ_1 3’) were directed to the nucleocapsid gene in region of conserved sequence between the variant viruses. Virus isolation/titration was also performed with the samples using Vero E6 cells using a standard SARS CoV-2 plaque assay as described ^35^.

Results were evaluated for statistical significance with GraphPad PRISM software. Mortality curves were evaluated using Mantel-Cox Log-Rank analysis. Average weight loss, clinical sign and virus titration data were compared with two-way ANOVA. Individual time points in particular assays were compared between two treatments with a two-tailed Student’s t test. Neutralization data was analyzed and PRNT_50_/PRNT_80_ calculated using Graphpad PRISM and the asymmetric sigmoidal 5PL standard curve fit (confidence limit 95%).

